# VARIETAL SCREENING OF BLACKGRAM GENOTYPES AGAINST CERCOSPORA LEAF SPOT (*Cercospora canescens*)

**DOI:** 10.1101/2024.08.12.607552

**Authors:** Siddhanta Shrestha, Sinchan Bohara, Sarita Oli, Anjal Nainabasti, Khem Raj Bohara, Kriti Upadhyaya, Indira Basnet

**Affiliations:** Institute of Agriculture and Animal Sciences, Tribhuvan University, Nepal; Department of Agronomy, Agriculture and Forestry University, Nepal

**Keywords:** AUDPC, Disease Scoring, Disease Severity, Legume, Blackgram, Varietal Screening

## Abstract

Screening of 12 blackgram genotypes against Cercospora Leaf Spot (*Cercospora canescens*) was carried out in RCB design with 3 replications in research plot of Mid-West Academy and Research Institute, Tulsipur, Dang during Bhadra to Mangsir, 2078 B.S. The blackgram genotypes were brought from Grain Legumes Research Program, Khajura, Banke. Disease severity was taken 3 times at 40, 47, and 54 days after sowing. Disease scoring was done as a percentage of leaf area infected on the individual plant at 7-day intervals and disease incidence, disease severity, mean AUDPC, and mean yield was calculated. Disease incidence was not significant among the tested genotypes. Disease severity at 40, 47, and 54 DAS was highly significant among the genotypes. Mean disease score and mean area under disease progressive curve (AUDPC) were also highly significant. Among the genotypes, 10 genotypes were categorized as moderately resistant and 2 genotypes (BLG 0066-1-1 and BLG 0035-1) were categorized as moderately susceptible. The highest Mean AUDPC value (324.1) was possessed by BLG 0035-1 followed by BLG 0066-1-1 (317.6). The lowest mean AUDPC value (175) was possessed by BLG 0069-1. A highly significant difference was found in yield among the black gram genotypes. The highest yield (799 kg/ha) by obtained by BLG 0068-2 followed by Rampur mas (769 kg/ha). The lowest yield (495 kg/ha) was obtained by BLG 0066-1.

## 1. Introduction

Blackgram (Vigna mungo L.), also known as urd bean or black matpe, is an important pulse crop belonging to the Fabaceae family. In Nepal, it is a popular legume dish referred to as “maas ko daal” or “kalo daal” and is commonly served with rice and curry. Blackgram cultivation in Nepal covers an area of 23,056 hectares, with a production of 20,440 metric tons and productivity of 887 kg/ha (MoAD, 2022).

Blackgram is a nutrient-rich crop that can thrive in diverse climatic conditions due to its ability to fix atmospheric nitrogen through root nodulation, thereby improving soil fertility and productivity. However, blackgram production in Nepal faces significant yield losses due to various biotic and abiotic constraints. Among the biotic factors, Cercospora leaf spot (CLS) disease, caused by the fungal pathogen *<u>Cercospora canescens</u>*, poses a severe threat to Vigna species cultivation in South Asian countries, including Nepal, where hot and humid conditions prevail (Gupta et al., 2014).

CLS was first reported in Delhi, India (Munjal et al., 1960), and has since spread to other parts of Asia and tropical regions worldwide (Pandey et al., 1983). The disease is highly prevalent in the humid and tropical areas of Nepal, India, Bangladesh, Indonesia, Malaysia, Philippines, and Thailand. Under natural epiphytotic conditions, CLS can cause yield losses of up to 96% in blackgram (Kaur et al., 2004). The fungus primarily infects Vigna species during the flowering stage, initially causing necrotic lesions on leaves, followed by spot development. During the pod-filling stage, the disease spreads rapidly, leading to premature defoliation, reduced plant size, and smaller leaves, pods, and seeds, resulting in yield losses ranging from 46% to 61% (Chand et al., 2015; Sompong et al., 2011).

The average yield of blackgram in the region remains relatively low due to various factors, including the use of poor-quality seeds, soil constraints, and pest and disease problems. Nepalese farmers often cultivate blackgram without proper scientific knowledge, land preparation, and crop management practices, leading to heavy infestation by pests and diseases. To control CLS, farmers rely on fungicides without considering their adverse effects, mode of action, spray formulation, and required waiting periods, potentially increasing health, ecological, and pathogenic risks.

Developing durable and effective disease resistance through the identification of resistant and high-yielding blackgram varieties is crucial for sustainable disease management. However, cultivars with a higher level of resistance to CLS are not readily available, and disease severity varies among varieties. Therefore, this research aims to identify resistant and susceptible blackgram genotypes to strengthen resistance breeding programs for blackgram in Nepal and contribute to the following objectives: a. to determine the incidence of Cercospora leaf spot disease; b. to evaluate disease severity among the tested blackgram genotypes; c. to estimate the mean area under the disease progress curve (AUDPC) and yield performance of the blackgram genotypes

## 2. Materials and Methods

### 2.1 Study Site

The experiment was conducted during the summer season of 2021 at the field site and pathology lab of the Institute of Agriculture and Animal Sciences (IAAS), MARICOLS, Dang district, Nepal. The location (27.9904°N, 82.3018°E, 725 m above mean sea level) falls within the inner Terai region of Nepal, with a subtropical climate.

### 2.2 Plant Materials

Twelve blackgram (Vigna mungo L.) genotypes, namely Rampur Mas, BLG 0068-2, BLG 0076-2, BLG 0069-1, BLG 0061-2-2, Khajura Mas-1, BLG 0066-1-1, BLG 0068-1-1, BLG 0072-1, BLG 0035-1, Shekhar, and BLG 041-1, were obtained from the Grain Legume Research Program, Khajura, Banke, Nepal.

### 2.3 Experimental Design and Layout

The experiment was laid out in a Randomized Complete Block Design (RCBD) with three replications. Each genotype was allocated to a 2 m × 1 m plot, with a row-to-row spacing of 25 cm. The gross experimental area was 84 m^2^, and the layout consisted of three rows, each containing 12 plots. Well-decomposed farmyard manure (FYM) at 15 kg/m^2^ was applied during field preparation, and inorganic fertilizers (urea, diammonium phosphate, and potash) were supplied at the recommended dose of 20:40:20 N:P:K kg/ha.

**Table 1:** List of blackgram genotypes used for the experiment.

| S.N. | Treatments | Symbol |
| --- | --- | --- |
| 1 | Rampur mas | T1 |
| 2 | BLG 0068-2 | T2 |
| 3 | BLG 0076-2 | T3 |
| 4 | BLG 0069-1 | T4 |
| 5 | BLG 0061-2-2 | T5 |
| 6 | Khajura mas-1 | T6 |
| 7 | BLG 0066-1-1 | T7 |
| 8 | BLG 0072-1 | T8 |
| 9 | BLG 0072-1 | T9 |
| 10 | BLG 0035-1 | T10 |
| 11 | Shekhar | T11 |
| 12 | BLG 0041-1 | T12 |

**Figure 1:**
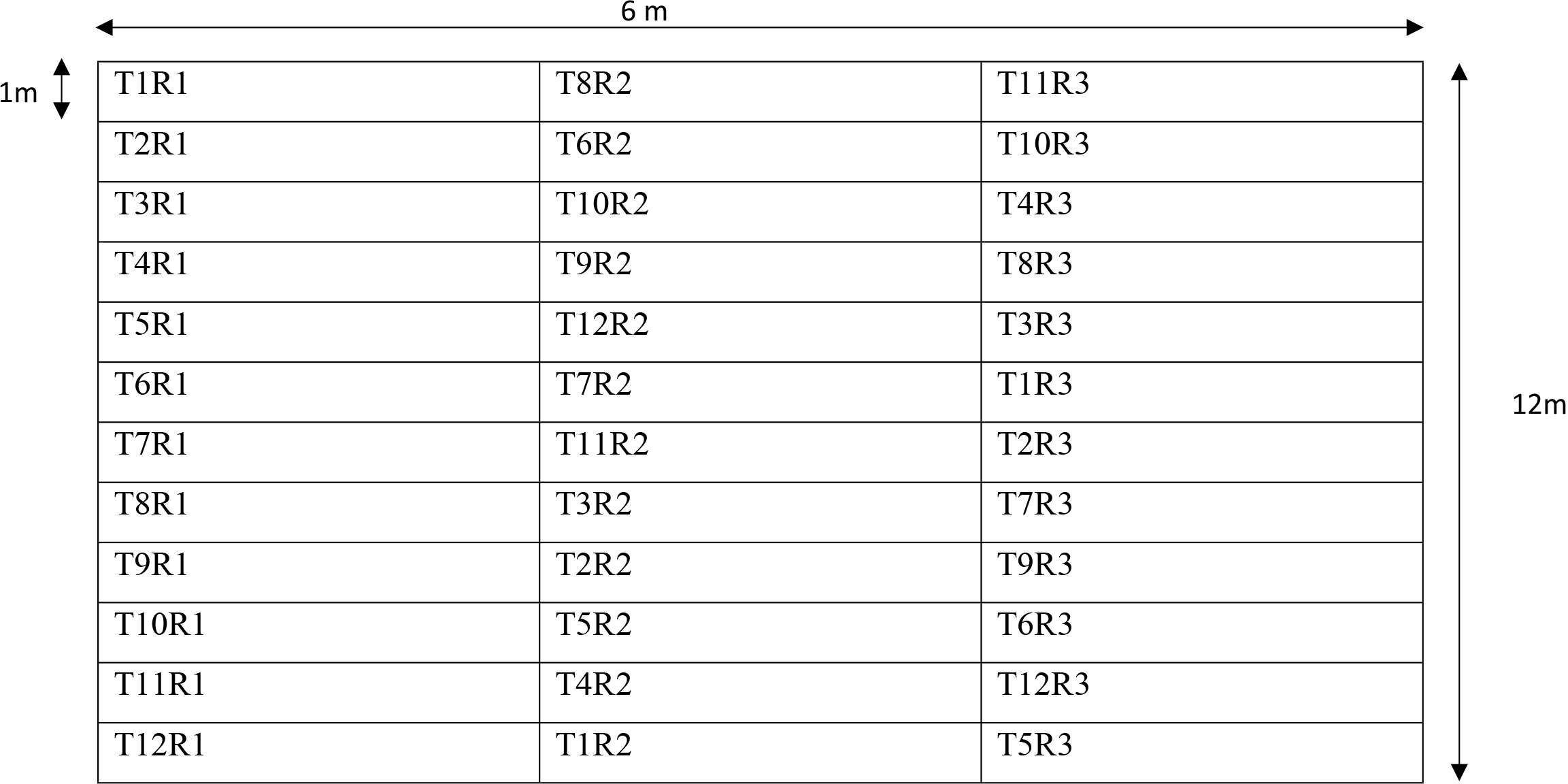
Layout of the experimental field during Bhadra, 2078.

#### Individual plot

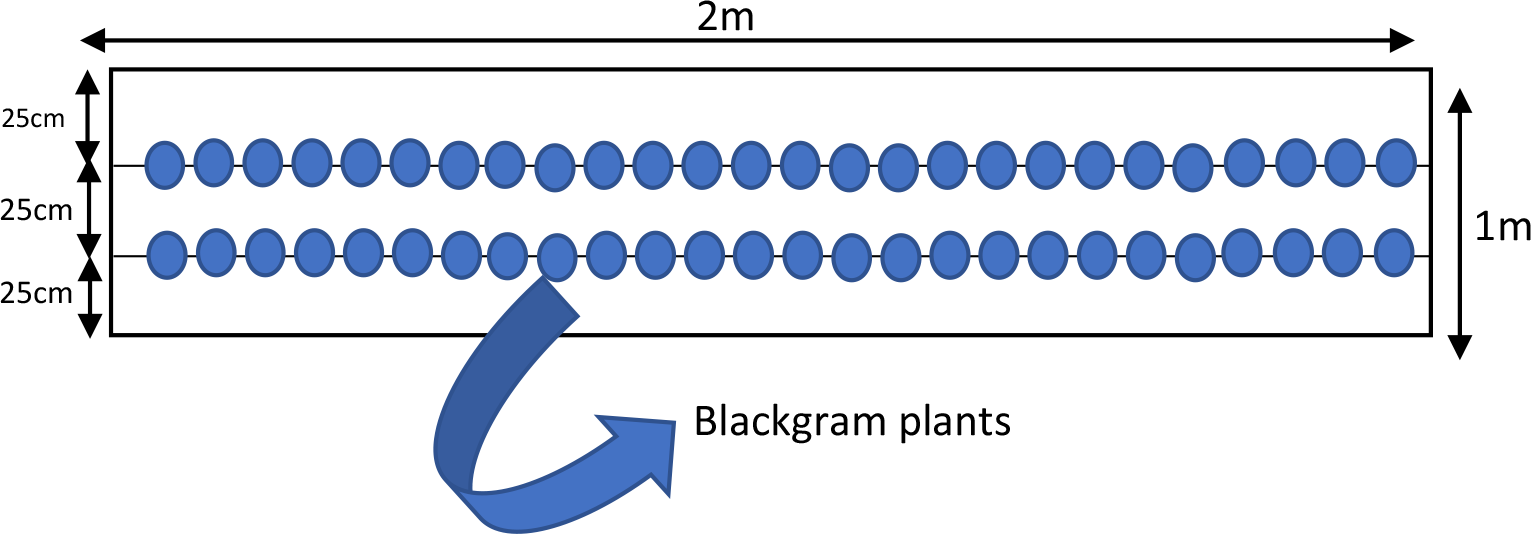

### 2.4 Cultural Practices

Land preparation was carried out on 26th Bhadra (September 11, 2021), involving three rounds of plowing, planking, and leveling. Raised beds of 10-15 cm height were prepared for sowing to avoid waterlogging. Seeds were sown on the raised beds by the line sowing method at a rate of 6 g per plot. Hand weeding was performed twice, on the 13th-14th and 39th-40th days after sowing (DAS). Chlorpyrifos and cypermethrin (2 ml/L) were sprayed on the 10th of Kartik (October 24, 2021) to control hairy caterpillars. No irrigation was provided due to sufficient rainfall during the crop growth period.

### 2.5 Disease Assessment

#### 2.5.1 Disease incidence

The number of germinated plants per plot of the individual genotypes was counted. After the appearance of the disease, disease plant was counted for computing CLS incidence Cercospora leaf spot disease incidence was calculated by the following;

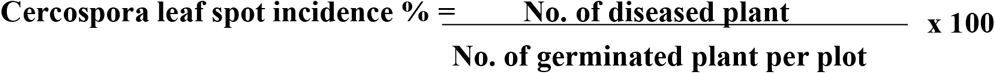

#### 2.5.2 Disease scoring

Disease incidence was recorded after the appearance of the disease (3 times scoring was done at the interval of 7 days at 40 DAS, 47 DAS, and 54 DAS). These scorings were taken at 7 days intervals during the pod formation stage. The data obtained from the experiment were grouped into five categories as resistant (R), moderately resistant (MR), moderately susceptible (MS), susceptible (S), and highly susceptible (HS) types to determine the nature of genotypes. The score 1 was considered as resistant whereas 2-3 was moderately resistant, 4-6 as moderately susceptible, 6 as susceptible, and 7-9 were considered highly susceptible. The disease scoring chart was given by (Alice & Nadarajan, 2007) is given as:

From the score taken, disease severity was calculated per plot by using the following formula:

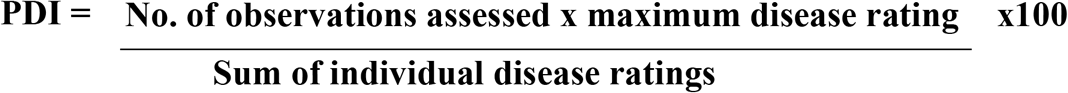

#### 2.5.3 Estimation of the area under disease progress curve

The area under the disease progress curve (AUDPC) was calculated by summarizing the progress of disease severity. The pattern of the epidemic in terms of the number of lesions, amount of diseased tissue, or the number of the diseased plant is given by a curve called disease progress curve, that shows epidemic over time and the area covered by this curve is known as AUDPC. AUDPC was computed, from the disease severities values from the formula given by (Shaner & Finney, 1977) (Das, Rajaram, Mundt, & Kronstad, 1992)

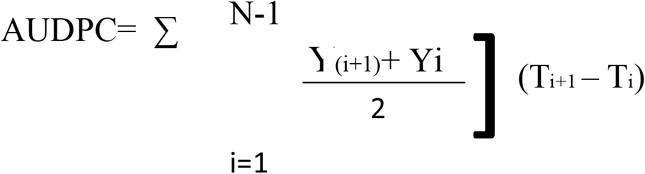

Where,

Yi= disease severity on the first date

Ti= date on which the disease was scored

N= number of dates on which disease was scored.

### 2.6 Statistical Analysis

Data were tabulated and analyzed using M-STAT software. Analysis of variance (ANOVA) was performed at a 5% level of significance to test for differences among genotypes for disease incidence, severity, AUDPC, and yield. Mean separation was conducted using the least significant difference (LSD) test. A regression analysis was performed to quantify the relationship between AUDPC and yield.

## 3. Results

### 3.1 Disease Incidence

Analysis of variance (ANOVA) revealed no significant differences (P = 0.66) in Cercospora leaf spot disease incidence among the 12 blackgram genotypes evaluated (Table 2). Disease incidence ranged from 6.3% in BLG 0066-1-1 to 10.3% in BLG 0035-1, with an overall mean of 7.7% and a coefficient of variation (CV) of 15%.

**Table 2:** Disease Score Chart.

| **Grade** | **Percentage Infection** | **Reaction** |
| --- | --- | --- |
| 1 | No infection on leaves | Resistant (R) |
| 2 | 0.1% to 5% infection on the leaf surface | Moderately resistant (MR) |
| 3 | 5.1% to 10% infection on the leaf surface | Moderately resistant (MR) |
| 4 | 10.1% to 15% infection on the leaf surface | Moderately susceptible (MS) |
| 5 | 15.1% to 30% infection on the leaf surface | Moderately susceptible (MS) |
| 6 | 30.1% to 40% infection on the leaf surface | Susceptible (S) |
| 7 | 40.1% to 50% infection on the leaf surface | Highly susceptible (HS) |
| 8 | 50.1% to 75% infection on the leaf surface | Highly susceptible (HS) |
| 9 | Above 75% infection on the leaf surface | Highly susceptible (HS) |

### 3.2 Disease Severity

Highly significant differences (P < 0.01) in disease severity were observed among genotypes at 40, 47, and 54 days after sowing (DAS) (Table 3). At 40 DAS, the genotype BLG 0035-1 exhibited the highest disease severity (40.74%), followed by BLG 0066-1-1 (37.04%), while BLG 0069-1 and Shekhar displayed the lowest severity (18.52%). The disease progressed, and at 47 DAS, BLG 0066-1-1 and BLG 0035-1 showed maximum severity (44.44%), with BLG 0068-2 and Shekhar exhibiting minimum values (25.93%). A similar trend continued at 54 DAS, where BLG 0066-1-1 and BLG 0035-1 had the highest disease severity (55.55%), and Shekhar remained the least affected genotype (25.93%).

**Table 3:** Cercospora leaf spot disease incidence of blackgram genotypes.

| S.N. | Treatments | Incidence % |
| --- | --- | --- |
| 1 | Rampur Mas | 6.8 |
| 2 | BLG 0068-2 | 7.6 |
| 3 | BLG 0076-2 | 6.8 |
| 4 | BLG 0069-1 | 6.5 |
| 5 | BLG 0061-2-2 | 7.6 |
| 6 | Khajura Mas-1 | 7.5 |
| 7 | BLG 0066-1-1 | 6.3 |
| 8 | BLG 0068-1-1 | 7.2 |
| 9 | BLG 0072-1 | 9.7 |
| 10 | BLG 0035-1 | 10.3 |
| 11 | Shekhar | 8.4 |
| 12 | BLG 041-1 | 6.6 |
| Probability |  | 0.66 |
| CV% |  | 15 |
| LSD |  | NS |

CV%: Coefficient of Variation, LSD: Least Significant Difference, NS: Non-significant

### 3.3 Disease Score, AUDPC, and Genotype Reaction

The genotypes differed significantly (P < 0.01) in their mean disease score and area under disease progression curve (AUDPC) values (Table 4). Mean disease scores ranged from 2.1 in Shekhar to 4.3 in BLG 0035-1, with of 9.79%. Likewise, mean AUDPC values varied between 168.5 for Shekhar and 324.1 for BLG 0035-1 (CV = 9.07%). Based on these parameters, the genotypes were categorized into different resistance groups: BLG 0066-1-1 and BLG 0035-1 were moderately susceptible, while the remaining ten genotypes, including Rampur Mas, BLG 0068-2, BLG 0076-2, BLG 0069-1, BLG 0061-2-2, Khajura Mas-1, BLG 0068-1-1, BLG 0072-1, Shekhar, and BLG 041-1, were moderately resistant to Cercospora leaf spot disease.

**Table 4:** Disease severity of Blackgram genotypes at different date of scoring.

| S.N. | Treatments | Disease severity |  |  |
| --- | --- | --- | --- | --- |
|  |  | 40 DAS | 47 DAS | 54 DAS |
| 1 | Rampur Mas | 22.22 cd | 33.33 bc | 29.63 de |
| 2 | BLG 0068-2 | 25.93 cd | 29.63 bc | 33.33 cde |
| 3 | BLG 0076-2 | 25.93 cd | 40.74 a | 44.44 b |
| 4 | BLG 0069-1 | 18.52 d | 25.93 c | 29.63 de |
| 5 | BLG 0061-2-2 | 25.93 cd | 33.33 bc | 40.74 bc |
| 6 | Khajura Mas-1 | 29.63 bc | 33.33 bc | 37.04 bcd |
| 7 | BLG 0066-1-1 | 37.04 ab | 44.44 a | 55.55 a |
| 8 | BLG 0068-1-1 | 29.63 bc | 33.33 bc | 37.04 bcd |
| 9 | BLG 0072-1 | 29.63 bc | 37.04 ab | 40.74 bc |
| 10 | BLG 0035-1 | 40.74 a | 44.44 a | 55.55 a |
| 11 | Shekhar | 18.52 d | 25.93 c | 25.93 e |
| 12 | BLG 041-1 | 29.63 bc | 33.33 bc | 40.74 bc |
| SEm± |  | 3.10 | 2.34 | 2.65 |
| CV% |  | 19.38 | 11.76 | 11.75 |
| LSD |  | 9.11** | 6.88** | 7.79** |

DAS: Days after sowing, CV%: Coefficient of Variation, LSD: Least Significant Difference Sem: Standard Error Mean, Means followed by the same letter in a column are not significantly different at 5% level of significance, \*\*: Highly Significant

### 3.4 Yield Performance

Yield differed significantly (P < 0.01) among blackgram genotypes, ranging from 495 kg/ha in BLG 0066-1-1 to 799 kg/ha in BLG 0068-2 (Table 5). The genotypes Rampur Mas (769 kg/ha), BLG 0069-1 (750 kg/ha), BLG 0068-1-1 (721 kg/ha), and Shekhar (750 kg/ha) also exhibited high yield potential. Regression analysis revealed a strong negative correlation (R^2 = 0.833) between mean AUDPC and yield (Figure 2), indicating that 83.3% of the variation in yield could be attributed to disease severity.

**Table 5:** Mean Disease score, Mean AUDPC, and disease reaction of blackgram genotypes to Cercospora leaf spot disease.

| S.N. | Genotypes | Mean Disease score | Mean AUDPC | Reaction |
| --- | --- | --- | --- | --- |
| 1 | Rampur Mas | 2.5 ab | 207.4 cd | Moderately Resistant (MR) |
| 2 | BLG 0068-2 | 2.6 ab | 207.4 cd | Moderately Resistant (MR) |
| 3 | BLG 0076-2 | 3.3 ab | 265.7 b | Moderately Resistant (MR) |
| 4 | BLG 0069-1 | 2.2 b | 175.0 d | Moderately Resistant (MR) |
| 5 | BLG 0061-2-2 | 3.0 ab | 233.3 bc | Moderately Resistant (MR) |
| 6 | Khajura Mas-1 | 3.0 ab | 233.3 bc | Moderately Resistant (MR) |
| 7 | BLG 0066-1-1 | 4.1 a | 317.6 a | Moderately Susceptible (MS) |
| 8 | BLG 0068-1-1 | 3.0 ab | 233.3 bc | Moderately Resistant (MR) |
| 9 | BLG 0072-1 | 3.2 ab | 252.8 b | Moderately Resistant (MR) |
| 10 | BLG 0035-1 | 4.3 a | 324.1 a | Moderately Susceptible (MS) |
| 11 | Shekhar | 2.1 b | 168.5 d | Moderately Resistant (MR) |
| 12 | BLG 041-1 | 3.1 ab | 239.8 bc | Moderately Resistant (MR) |
| SEm± |  | 0.54 | 2.34 |  |
| CV% |  | 9.79 | 9.07 |  |
| LSD |  | 1.60 ** | 6.88** |  |

AUDPC: Area Under Disease Progress Curve

### 3.5 Regression Study

A regression analysis was performed to investigate the relationship between disease severity, represented by mean AUDPC values, and yield (Figure 2). The analysis revealed a strong negative correlation (R^2 = 0.833) between mean AUDPC and yield, indicating that 83.3% of the variation in yield could be attributed to disease severity. The remaining 16.7% of the yield variation was likely influenced by other factors not accounted for in this study. The negative correlation suggests that as disease severity increased, yield declined proportionally across the evaluated blackgram genotypes

**Table 6:**
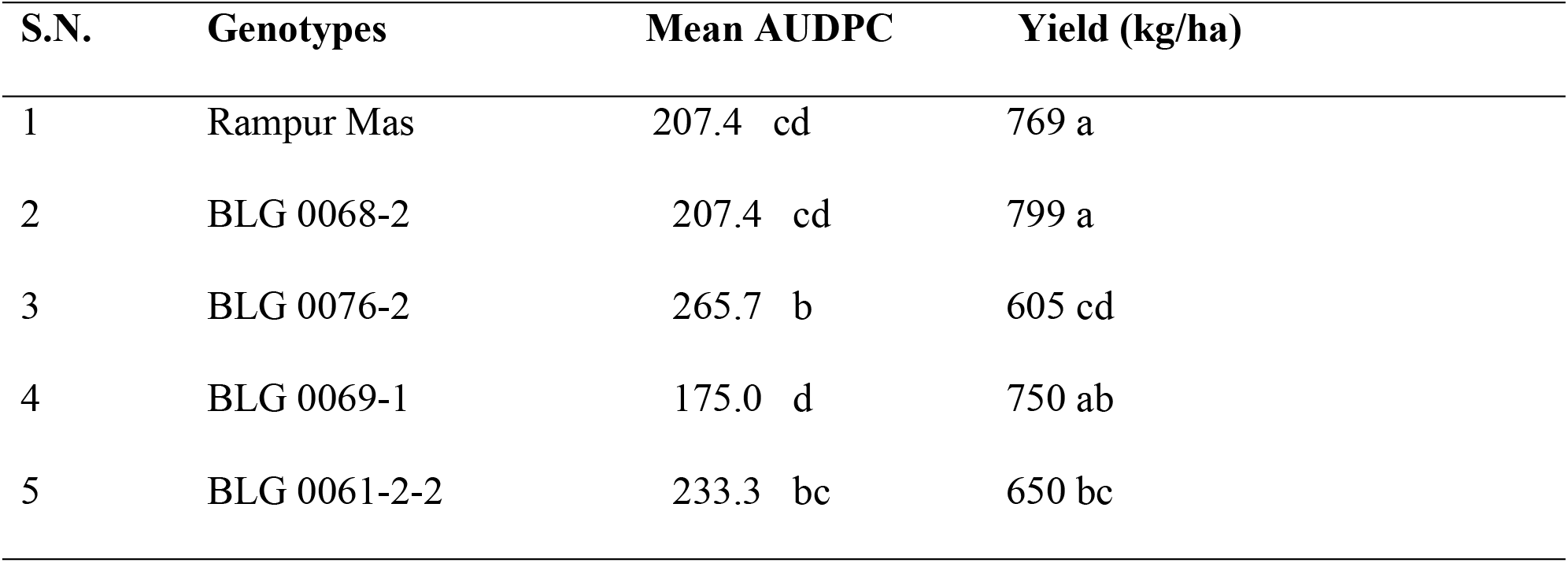
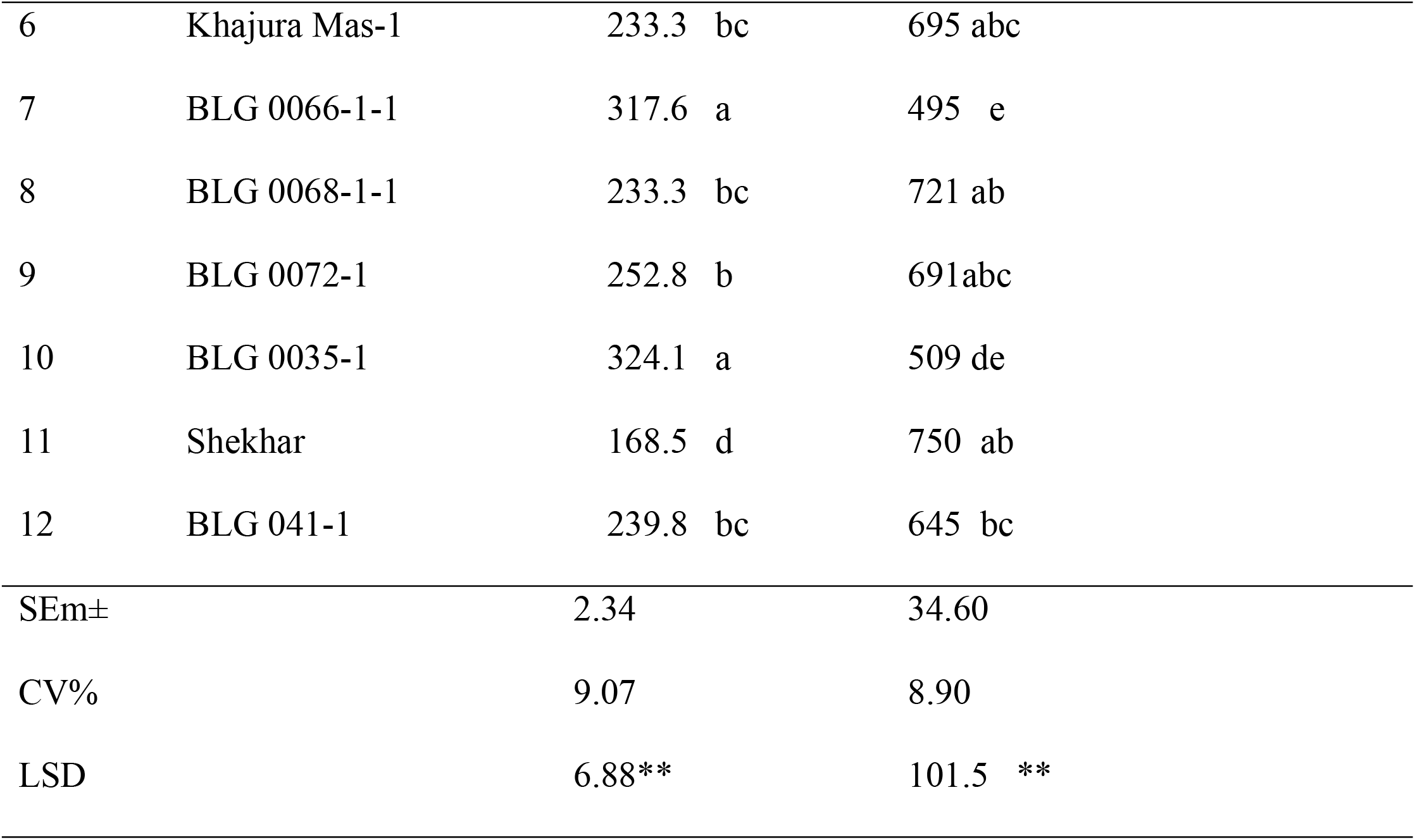
Mean AUDPC and yield of Blackgram genotypes.

| S.N. | Genotypes | Mean AUDPC | Yield (kg/ha) |
| --- | --- | --- | --- |
| 1 | Rampur Mas | 207.4 cd | 769 a |
| 2 | BLG 0068-2 | 207.4 cd | 799 a |
| 3 | BLG 0076-2 | 265.7 b | 605 cd |
| 4 | BLG 0069-1 | 175.0 d | 750 ab |
| 5 | BLG 0061-2-2 | 233.3 bc | 650 bc |
| 6 | Khajura Mas-1 | 233.3 bc | 695 abc |
| 7 | BLG 0066-1-1 | 317.6 a | 495 e |
| 8 | BLG 0068-1-1 | 233.3 bc | 721 ab |
| 9 | BLG 0072-1 | 252.8 b | 691abc |
| 10 | BLG 0035-1 | 324.1 a | 509 de |
| 11 | Shekhar | 168.5 d | 750 ab |
| 12 | BLG 041-1 | 239.8 bc | 645 bc |
| SEm± |  | 2.34 | 34.60 |
| CV% |  | 9.07 | 8.90 |
| LSD |  | 6.88** | 101.5 ** |

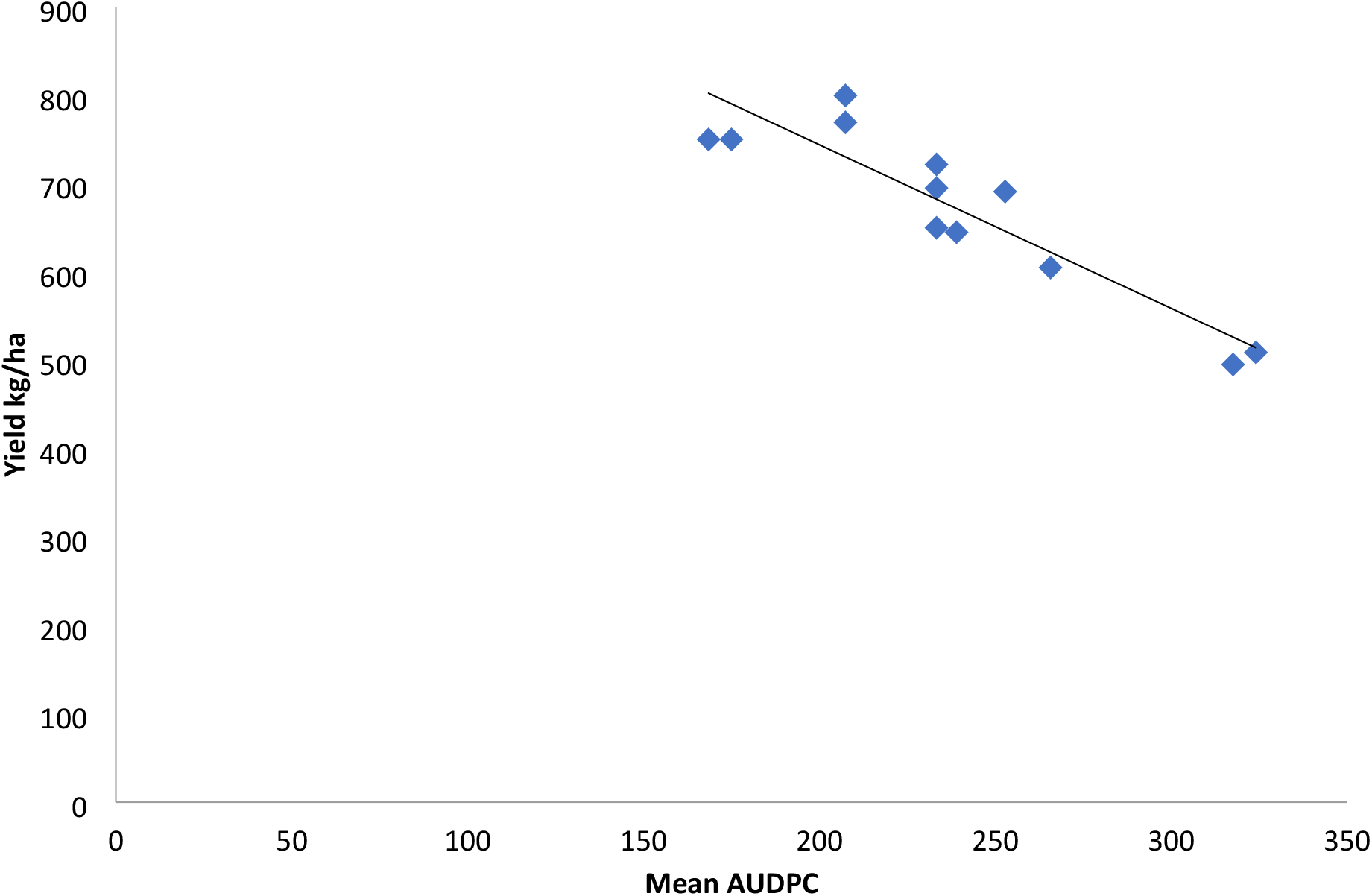

## 4. Discussion

The results of this study provide valuable insights into the resistance of various blackgram genotypes to Cercospora leaf spot disease. The analysis of variance revealed no significant differences in disease incidence among the 12 genotypes evaluated, with incidence ranging from 6.3% to 10.3%. This suggests that all genotypes exhibited similar levels of resistance to initial infection by the pathogen(Bhaskar, 2017; Nair et al., 2024).

However, significant differences were observed in disease severity at 40, 47, and 54 days after sowing (DAS). At 40 DAS, BLG 0035-1 and BLG 0066-1-1 showed the highest disease severity, while BLG 0069-1 and Shekhar were the least affected. This trend continued at 47 and 54 DAS, with BLG 0035-1 and BLG 0066-1-1 consistently exhibiting the highest severity and Shekhar remaining the least affected genotype. These findings indicate that while initial infection rates were similar, some genotypes were more susceptible to disease progression and development of symptoms over time.

The mean disease score and area under disease progression curve (AUDPC) values also differed significantly among genotypes. BLG 0035-1 and BLG 0066-1-1 had the highest mean disease scores and AUDPC values, categorizing them as moderately susceptible. The remaining ten genotypes, including Rampur Mas, BLG 0068-2, BLG 0069-1, and Shekhar, were classified as moderately resistant. These results corroborate the findings of Adhikari et al., (2014), who evaluated the resistance of various blackgram genotypes to Cercospora leaf spot and identified similar levels of resistance.

Yield also differed significantly among genotypes, ranging from 495 kg/ha in BLG 0066-1-1 to 799 kg/ha in BLG 0068-2. Regression analysis revealed a strong negative correlation (R^2 = 0.833) between mean AUDPC and yield, indicating that 83.3% of the variation in yield could be attributed to disease severity. This finding is consistent with the study by Bhardwaj & Thakur, (2000), who reported a significant reduction in yield due to Cercospora leaf spot disease in blackgram.

The results of this study have important implications for breeding programs aimed at developing blackgram varieties with improved resistance to Cercospora leaf spot disease. The identification of moderately resistant genotypes, such as Rampur Mas, BLG 0068-2, BLG 0069-1, and Shekhar, provides valuable sources of resistance that can be utilized in breeding efforts. Additionally, the strong negative correlation between disease severity and yield highlights the importance of selecting for resistance to minimize yield losses in blackgram production.

## 5. Conclusion

Cercospora leaf spot, incited by *<u>Cercospora canescens</u>*, is a major biotic constraint for blackgram (Vigna mungo) cultivation in Nepal, causing substantial yield losses in susceptible varieties. This study evaluated twelve blackgram genotypes for resistance against the disease under field conditions at Tulsipur, Dang, during the 2078 B.S. season. Based on the area under the disease progress curve (AUDPC), none of the genotypes exhibited complete resistance, while BLG 0066-1-1 and BLG 0035-1 were categorized as moderately susceptible. The remaining ten genotypes, including Rampur Mas and BLG 0068-2, were classified as moderately resistant. Although no completely resistant source was identified, the moderately resistant genotypes represent valuable genetic resources for resistance breeding programs aimed at developing high-yielding, disease-resistant blackgram varieties. Continuous multi-environment trials are recommended to validate these findings and identify potential sources of complete resistance, contributing to sustainable blackgram productivity and enhanced food security in Nepal.

